# RNA length has a non-trivial effect in the stability of biomolecular condensates formed by RNA-binding proteins

**DOI:** 10.1101/2021.10.07.463486

**Authors:** Ignacio Sanchez-Burgos, Jorge R. Espinosa, Jerelle A. Joseph, Rosana Collepardo-Guevara

## Abstract

Biomolecular condensates formed via liquid–liquid phase separation (LLPS) play a crucial role in the spatiotemporal organization of the cell material. Nucleic acids can act as critical modulators in the stability of these protein condensates. Here, we present a multiscale computational strategy, exploiting the advantages of both a sequence-dependent coarse-grained representation of proteins and a minimal coarse-grained model that represent proteins as patchy colloids, to unveil the role of RNA length in regulating the stability of RNA-binding protein (RBP) condensates. We find that for a constant RNA/protein ratio in which phase separation is enhanced, the protein fused in sarcoma (FUS), which can phase separate on its own—i.e., via homotypic interactions—only exhibits a mild dependency on the RNA strand length, whereas, the 25-repeat proline-arginine peptide (PR_25_), which does not undergo LLPS on its own at physiological conditions but instead exhibits complex coacervation with RNA—i.e., via heterotypic interactions—shows a strong dependence on the length of the added RNA chains. Our minimal patchy particle simulations, where we recapitulate the modulation of homotypic protein LLPS and complex coacervation by RNA length, suggest that the strikingly different effect of RNA length on homotypic LLPS versus complex coacervation is general. Phase separation is RNA-length dependent as long as the relative contribution of heterotypic interactions sustaining LLPS is comparable or higher than that committed by protein homotypic interactions. Taken together, our results contribute to illuminate the intricate physicochemical mechanisms that influence the stability of RBP condensates through RNA inclusion.

## I. INTRODUCTION

Cells require precise compartmentalization of their material into different organelles in order to function. While some of these organelles and compartments are shaped by physical membranes, many others are sustained by a mechanism called liquid–liquid phase separation (LLPS) [1–4]. Like oil and water, biomolecules including multivalent proteins and, in some cases, nucleic acids, can spontaneously demix into phase-separated droplets known as biomolecular condensates [5, 6]. Beyond compartmentalization, numerous vital roles have been recently associated with biomolecular condensates, including cell signaling [2, 7], formation of super-enhancers [8], genome organization [9–12], and aiding cells to sense and react to environmental changes [13] among many others [14–17]. Within the extensive class of biomolecules that can undergo phase separation at physiological conditions, RNA-binding proteins (RBPs) such as FUS [18–20], hnRNPA1 [21, 22], TDP-43 [23–25], TAF-15 [26, 27], G3BP1 [28, 29] or EWSR1 [26, 27, 30], have been widely investigated due to their implications in the stability of stress granules [31, 32], P granules [1, 33, 34] or RNA granules/bodies [35–37]–important phase-separated organelles within cells.

Phase-separation of RBPs can be both promoted or inhibited by the presence of RNA in an RNA-concentration, and sometimes RNA-structure, dependent manner [27, 30, 38–48]. From the physicochemcial point of view, RBPs possess key features that explain their highly RNA–sensitive phase behaviour; i.e., they are multidomain proteins that combine aromatic-rich and arginine-rich intrinsically disordered regions (IDRs) [26, 49] – boosting the RBP’s multivalency needed for LLPS – with globular domains that exhibit high affinity for RNA (termed RNA recognition motifs (RRMs)) [50]. Hence, RPBs and RNA can establish both specific RNA-RRM interactions and non-specific electrostatic, cation-*π* and *π*-*π* interactions. To gain a mechanistic understanding of the intricate modulation of RBP condensate stability by RNA, experiments where single amino acids are mutated and/or post-translationally modified (e.g. phosphorylated [10, 11, 51] or methylated [18, 52, 53]) are of great value. Alongside, sequence-dependent molecular simulations can help uncover how specific protein regions, amino acid-RNA interactions, or RNA properties influence the experimentally observed behavior [47, 48, 54–56].

Computer simulations have been instrumental in advancing the characterization of biomolecular condensates from a thermodynamic, molecular and mechanistic perspective [6, 57–59]. Many approaches, such as atomistic Molecular Dynamics (MD) simulations [59–61], sequence-dependent high-resolution coarse-grained models [54, 62–65] or minimal representations of proteins [66–70], as well as lattice-based simulations [71–74] and mean field models [75–79] have been developed and exploited to interrogate biomolecular LLPS. These approaches have proved extremely useful for rationalizing the effects of key factors in LLPS, encompassing protein length [80, 81], amino acid sequence [54, 62, 63, 82, 83], multivalency [71, 84–90], conformational flexibility [91, 92], multicomponent composition [47, 67, 93–97], and elucidating the links between chemical modifications, sequence mutations, and protein–protein or protein–DNA interactions [98–103]. Moreover, coarse-grained models have been employed to investigate the RNA-induced reentrant LLPS behaviour of RBPs [47, 48], the effect of RNA on phase separation of small prion-like domains such as those of FUS, [67, 104], protamine [105] and LAF-1 [47], and the emergence of multiphasic protein–RNA condensates [106].

In this work, we seek to address another pertinent open question on the role of RNA in RBP LLPS [107]: What is the function of RNA strand length in biomolecular LLPS? For this, we use our recently developed residue/nucleotide-resolution coarse-grained protein/RNA model [55], which predicts biomolecular phase diagrams in quantitative agreement with experiments. We demonstrate striking and contrasting effects of RNA length on the phase behaviour of RBPs. For RBPs like FUS, which can undergo LLPS via homotypic protein–protein interactions, low-to-moderate RNA concentrations invariably lead to moderate enhancement of condensate stability irrespective of the RNA length (for a fixed total nucleotide/protein concentration). In contrast, for RBPs like PR_25_ that undergo RNA-dependent complex coacervation (i.e., LLPS driven by heterotypic protein–RNA interactions), increasing RNA length at constant total nucleotide concentration significantly promotes condensate stability. Next, we use minimal coarse-grained simulations to look at the problem from a soft condensed matter perspective. Our minimal simulations reveal that the striking differences in the impact of RNA length on complex coacervation versus homotypic LLPS originates in the diversity of intermolecular connections that biomolecules employ in the different scenarios to sustain the condensed liquid-network of the condensates.

## II. RESULTS AND DISCUSSION

### A. Multiscale modelling approach for RBP–RNA phase separation

Biomolecular LLPS entails the self-assembly of thousands of different proteins and other biomolecules into liquid-like condensates; hence, these condensates are often not amenable to atomistic-level simulations. Instead, coarse-grained models including mean field simulations [75–79, 108], lattice-based models [71–74], and high-resolution sequence-dependent approaches [54, 62–65, 109], are becoming the go-to simulation methods for characterizing the mechanistic and molecular details of biomolecular condensates. Here, we employ two protein/RNA coarse-grained models of different resolutions, previously developed by us, to elucidate the role of RNA length in modulating LLPS of RBPs: (1) the Mpipi sequence-dependent residue-resolution coarse-grained force field for proteins and RNA [55], and (2) a minimal model in which proteins are represented as patchy particles, and RNA as self-repulsive flexible polymers [67, 88] (Figure 1 (a)).

**Figure 1:**
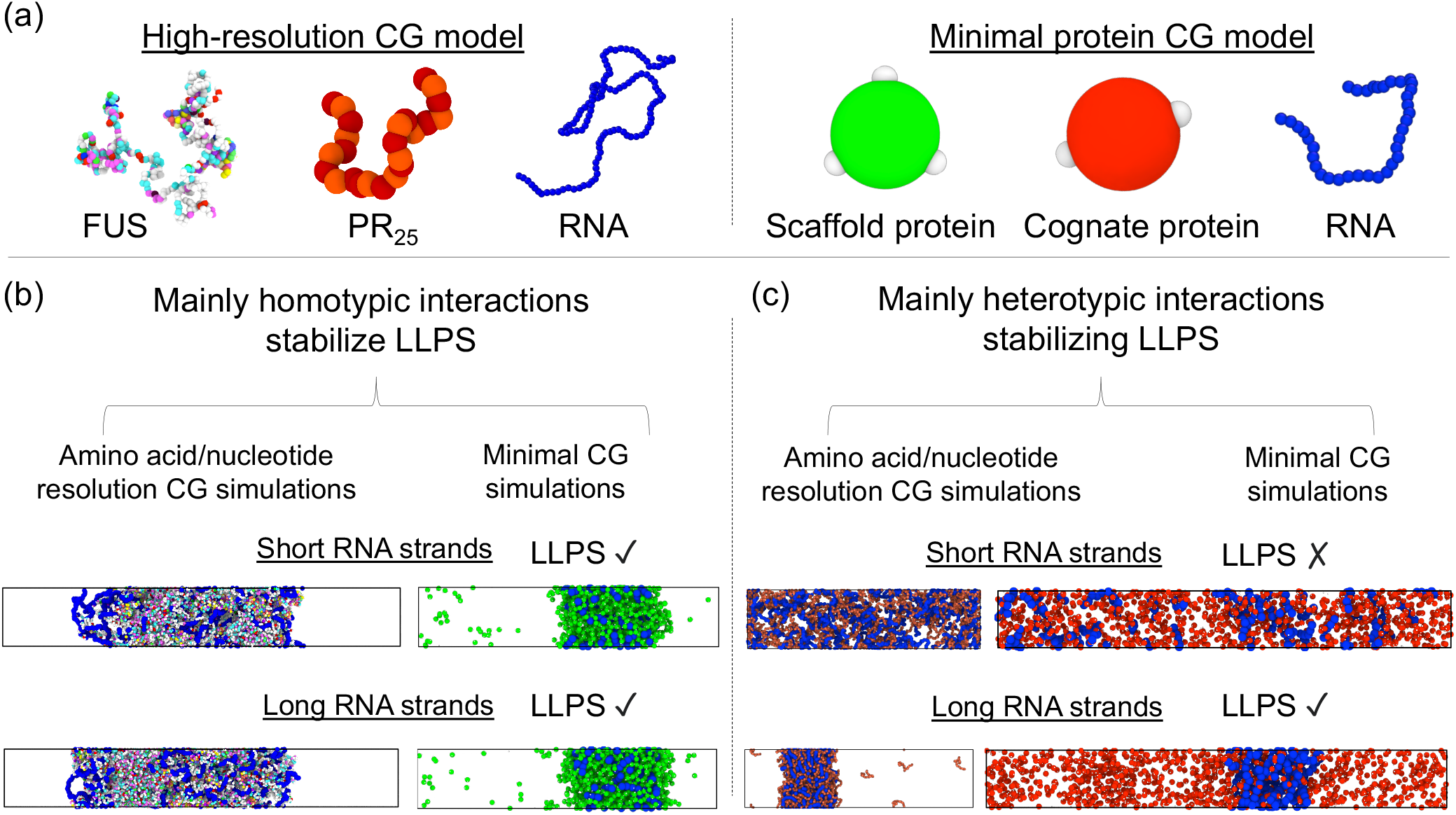
Coarse-grained models for investigating RBP–RNA phase-separation. (a) Left: High-resolution sequence-dependent representation of FUS, PR_25_ and a 400-mer polyU RNA strand. In this model [55], each amino acid and nucleotide is represented by a single bead with their own chemical identity while the solvent is implicitly described (please note that the size of the beads represented in panel (a) have been rescaled at convenience for visualization purposes). Globular protein domains are modelled as rigid-bodies regions based on the crystal structure of the folded domains, whereas IDR regions and RNA are fully flexible polymers. Coloured beads indicate distinct residues/nucleotides. (a) Right: Minimal model for scaffold proteins, cognate proteins and RNA [67]. White patches represent protein binding sites, while green and red spheres account for the excluded volume of the scaffold and cognate proteins respectively [88]. RNA is modelled via a self-repulsive fully flexible polymer of (pseudo) hard-spheres [67]. Please note that the real size of the RNA beads has been intentionally reduced to improve their visualization, and that their size is the same as the central pseudo hard-sphere of the proteins. (b) Direct Coexistence simulations of FUS/RNA (left) and scaffold proteins/RNA (right) using short RNA strands (top; 50-mer polyU and 10-bead RNA chains for FUS and the minimal scaffold protein model respectively) and long RNA strands (bottom; 400-mer and 250-bead RNA chains for FUS and the minimal scaffold protein model respectively) at *T/T*_*c*_=1.01, where *T*_*c*_ corresponds to the pure protein critical temperature of each system. (c) DC simulations of PR_25_ with RNA (left) and cognate proteins with RNA (right) using both short RNA strands (top; 40-mer polyU and 10-bead polyU RNA chains for PR_25_ and RNA cognate proteins respectively) and long RNA strands (bottom; 400-mer and 250-bead RNA chains for PR_25_ and RNA cognate proteins respectively) at *T/T*_*c*_=1.01, where *T*_*c*_ corresponds to the pure critical temperature of FUS (left) and scaffold proteins (right), as in panel (b).

Within the Mpipi model, protein residues and RNA bases are represented by single beads with unique chemical identities (Figure 1 (a) Left) in which hydrophobic, *π*–*π* and cation–*π* interactions are modelled through a mid-range pairwise potential (Wang-Frenkel potential [110]), and electrostatic interactions via Yukawa long-range potentials [54]. Bonded interactions between sequential amino acids within the same protein or nucleotides within the same RNA strand are described with a harmonic potential. Additionally, within Mpipi, the intrinsically disordered regions of the proteins and RNA strands are treated as fully flexible polymers, while globular domains are described as rigid bodies based on their corresponding crystal structures taken from the Protein Data Bank (PDB) and adapted to the model resolution. In the Mpipi model, the interactions between ‘buried’ amino acids within globular domains are scale down as proposed in Refs. [100, 111]. The physiological concentration of monovalent ions in solution (i.e., ∼150 mM NaCl), within the implicit solvent model is approximated by the screening length of the Yukawa/Debye-Hückel potential. Further details on the model parameters, protein sequences and simulation setups are provided in the Supplementary Material.

Complementary to the high-resolution sequence-dependent model, we employ a minimal coarse-grained model [67, 88] to investigate the role of RNA length in RBPs LLPS. Within this model, proteins are described by pseudo hard-sphere (PHS) [112] particles decorated with sticky patches that describe the protein binding sites (modelled through square-well-like potentials [113]); these allow the minimal proteins to establish multivalent transient interactions (Figure 1 (a) Right)). Additionally, RNA strands are modelled as fully flexible self-repulsive PHS polymers that can interact attractively with RBPs via mid-range non-specific interactions (see Supplementary Material and Ref. [67] for further details on the model potentials and parameters). Each minimal RNA bead accounts for tens of nucleotides and has the same size as the protein beads [67]. As in the residue-resolution coarse-grained model, an implicit solvent is used; accordingly, the diluted phase (i.e., the protein-poor liquid phase) and the condensed phase (i.e., the protein-rich liquid phase) are effectively a vapor and a liquid phase, respectively.

To measure the stability of the RBP–RNA condensates, we compute phase diagrams of the different systems in the space of temperature versus density by means of Direct Coexistence (DC) simulations [114, 115]. Within this approach, the two coexisting phases of the system are placed in the same simulation box; in our case, a high-density protein liquid and a very low-density one. We employ a rectangular box, with an elongated side perpendicular to the interfaces (long enough to capture the bulk density of each phase), while the parallel sides are chosen such that proteins cannot interact with themselves across the periodic boundaries [48]. We then run *NV T* MD simulations until equilibrium is reached. Once the simulations have converged, we measure the equilibrium coexisting densities of both phases along the long side of the box, excluding the fluctuations of the interfaces and keeping the center of mass of the system fixed. We repeat this procedure at different temperatures until we reach supercritical temperatures, where no phase separation is observed any longer. Then, to avoid finite system-size effects close to the critical point, we evaluate the critical temperature (*T*_*c*_) and density (*ρ*_*c*_) using the law of critical exponents and rectilinear diameters [116] (as shown in Refs. [67, 88]). Figure 1 (b) (Middle and Bottom panels) depicts phase-separated systems computed via DC simulations, while Figure 1 (c) (Middle panel) shows supercrticial systems (i.e., no phase separation)

### B. Impact of RNA length in the phase behaviour of FUS *versus* PR_25_ condensates

Using MD simulations of our protein/RNA sequence-specific Mpipi model [55], we first investigate the effect of adding disordered Poly-U single-stranded RNA chains to RBPs condensates, and varying the length of the Poly-U (while keeping the total amount of U nucleotides and protein constant). Specifically, we compare the effects of RNA-length in the phase behaviour of two different RBPs: (1) FUS, which can phase separate on its own at physiological conditions via homotypic protein–protein interactions, and (2) PR_25_, which undergoes LLPS at physiological conditions only in the presence of RNA via heterotypic RNA–protein interactions [46, 100, 111].

For the different FUS/RNA systems, regardless of the length of the RNA strands in each case, we always add a total amount of U nucleotides to get a constant U/FUS mass ratio of 0.38; this is because this ratio enhances phase separation with respect to the pure FUS system. Specifically, we test six polyU lenghts: (i) 64 polyU chains of 25 nucleotides each, (ii) 32 polyU chains of 50 nucleotides each, (iii) 16 polyU chains of 100 nucleotides each, (iv) 8 polyU chains of 200 nucleotides each, (v) 4 polyU chains of 400 nucleotides each, and (vi) 2 polyU chains of 800 nucleotides each. In all these systems (Figure 2 (a)), we observe a moderate increase in the critical temperature of FUS when RNA is added, independently of the length of RNA; i.e., all FUS+polyU systems we simulate have very similar critical temperatures within the uncertainty.

**Figure 2:**
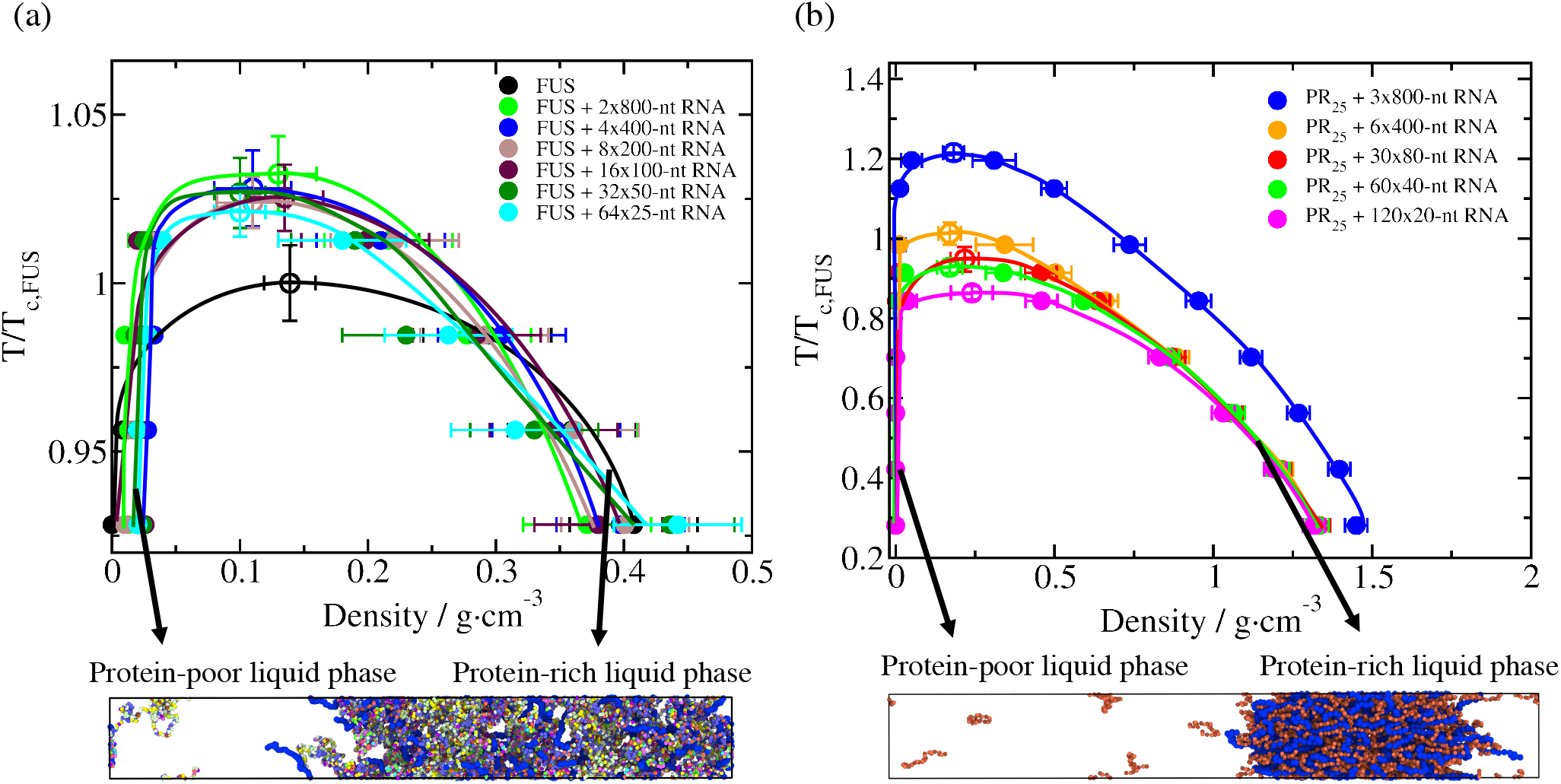
(a) Temperature–density phase diagrams of FUS with polyU RNA of different lengths at a constant polyU/FUS mass ratio of 0.38, and for a pure system of FUS (black curve). The length of polyU RNA strands range from 25-nucleotide to 800-nucleotide. (b) Temperature–density phase diagrams of PR_25_ with RNA at different lengths at a constant RNA/PR_25_ mass ratio of 1.36. RNA lengths range from 20-nucleotide to 800-nucleotide strands. In both (a) and (b) panels, filled circles represent the coexisting densities evaluated from DC simulations while empty circles depict the critical temperatures estimated from the law of rectilinear diameters and critical exponents [116] near the critical temperature. Temperature in both panels has been normalized by the critical temperature of pure FUS, *T*_*c,F US*_=355 K (black empty circle in (a)). Representative snapshots of the DC simulations used to compute the phase diagrams of both systems for a given RNA strand length (a) FUS-polyU(4×400-nt) and b) PR_25_-polyU(6×400-nt)) under phase-separating conditions are included below. The same color code employed in Fig. 1 applies here.

To determine if proteins that phase separate by complex coacervation exhibit a similar trend, we next investigate the effect of RNA length on PR_25_–polyU mixtures using Mpipi. In this case, we fix the polyU/PR_25_ mass ratio to 1.36, as this maximizes the size of the coexistence region for the smallest length of polyU used (20 nucleotides). We then test five different polyU lengths: (i) 120 polyU chains of 20 nucleotides each, (ii) 60 polyU chains of 40 nucleotides each, (iii) 30 polyU chains of 80 nucleotides each, (iv) 6 polyU chains of 400 nucleotides each, (v) 3 polyU chains of 800 nucleotides each. The dependence of the phase behaviour of PR_25_ on RNA length is strikingly different (Figure 2 (b)): the size of the coexistence region for PR_25_+polyU now grows continuously as the length of polyU increases. Indeed, lengthening RNA from 20 to 800 nucleotides increases the critical temperature by as much as 50%. This observation is significant, since while increasing the RNA length, we have maintained a constant nucleotide concentration, which ensures that the total number of binding sites in the RNA molecules available for protein binding, is the same in all cases.

To elucidate the molecular origin of this important difference, we compute the percentage of LLPS-stabilizing contacts per unit of volume at 350 K (*T/T*_c,FUS_ ∼ 1) for FUS (Figure 3 (a)), and 300 K in the case of PR_25_ (*T/T*_c,FUS_ ∼0.85 (Figure 3 (b)); note that PR_25_ cannot phase separate on its own). These temperatures were chosen as the highest temperatures at which phase separation is observed for each protein at all RNA lengths. We find that FUS+polyU condensates are mostly stabilized by protein–protein interactions, and more modestly contributed by protein–RNA interactions (Figure 3 (a)). This result suggests that within FUS condensates, where FUS acts as the scaffold, a moderate concentration of RNA creates a few more bridges among the scaffolds—acting as molecule that increases the effective valency of FUS within the condensate or co-scaffold in phase separation [117, 118]. We also note that increasing the concentration of polyU in our FUS-poly-U mixtures, rather than the RNA length at constant U/FUS ration, would eventually inhibit FUS phase separation (the so-called RNA-induced LLPS reentrant behaviour [27, 40, 45–48]). At physiological conditions, FUS–FUS interactions are sufficient to drive the system to phase separate [119]. Addition of a moderate amount of RNA creates more connections between FUS proteins by directly binding to free sites on FUS [27] (especially via specific RNA–RRM interactions and non-selective electrostatic and *π*-*π* interactions [30, 38–44, 48]), while high amounts of RNA begin to outcompete the FUS–FUS connections and introduce electrostatic repulsion that eventually inhibit LLPS. At moderate concentrations, RNA marginally increases the connectivity of an already sufficiently connected condensed liquid network [46]. This is evident from the density of FUS–FUS and FUS–RNA contacts remaining almost constant as the length of the RNA strands increases (Figure 3 (a)), following the same trend of critical points as a function of RNA length in the mixtures (Figure 3 (c)). We reason that RNA length does not have a strong impact in the stability of FUS condensates because: (1) the total number of FUS–RNA bonds is low enough that the competition between RNA–RNA repulsion among short RNA chains (that would be reduced by the covalent bonds among longer RNA chains) and RNA–FUS attraction becomes unimportant, and (2) FUS is a large protein that offers many distant RNA-binding sites that are equally viable for moderately short RNA chains that repel each other, or for long RNA chains that are stitched together by covalent bonds, as long they have a comparable radius of gyration to that of the proteins [48]. Despite this, we note that experiments have reported how RNA length can modulate the stability of some RNA-binding proteins such as FUS [120] or LAF-1 [121]. However, in those cases the difference in stability was observed at very short lengths (i.e., ∼20-40 nucleotides), where the RNA strands were much smaller than the proteins themselves. In fact, when RNA length is not long enough to bind to more than one protein at the time, it can hinder the association with other proteins as recently shown in Ref. [48]. Our results argue that as long as the length of the RNA strands is sufficient to allow single RNA molecules to simultaneously bind to more than one RBP at a time (Fig. 3 (a)), the effect of RNA length on RBPs homotypic phase separation is expected to be marginal (Fig. 3 (c)).

**Figure 3:**
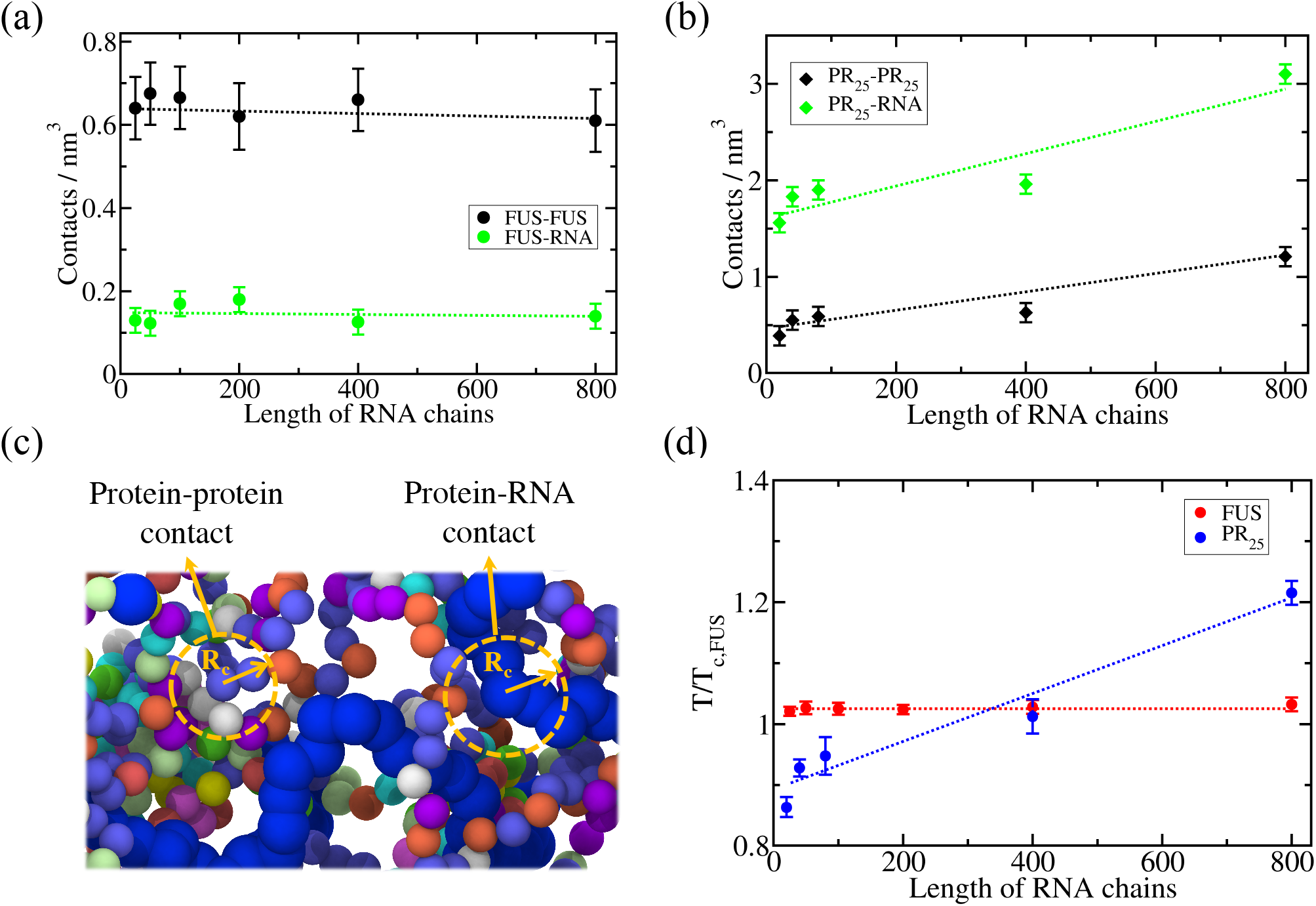
Density of LLPS-stabilizing intermolecular contacts within condensates as a function of RNA length plotted separately for protein–protein interactions (black circles) and protein–RNA interactions (green circles) for FUS–polyU (a) and PR_25_-polyU mixtures (b). The temperature at which the intermolecular contacts were computed was *T/T*_c,FUS_=1 for FUS–RNA systems and *T/T*_c,FUS_=0.85 for PR_25_–RNA mixtures. (c) Representative snapshot of a bulk FUS-polyU condensate to illustrate the employed cut-off distance (*R*_*c*_) criterion to identify protein-protein and protein-RNA contacts. The same color code described in Fig. 1 applies here. (d) Critical temperature *versus* RNA length for FUS–RNA (red) and PR_25_–RNA (blue) systems. The RNA/protein mass ratio of all systems was kept constant at 0.38 for FUS–RNA systems and at 1.36 for PR_25_–RNA mixtures.

In contrast, PR_25_ condensates are mostly stabilized by polyU–PR_25_ interactions and only modestly by protein– protein interactions (Figure 3 (b)), as expected from their complex coacervation being dependent on the presence of polyU. Furthermore, consistent with the increase of the critical temperature with RNA length (Figure 3 (c)), the density of protein–RNA intermolecular contacts increases significantly as the RNA lengthens, especially at chain lengths of hundreds of nucleotides (i.e., 800-mer polyU chains in our simulations). Because PR_25_ must bind to RNA to form a liquid network, adding covalent bonds within the RNA chains—-for instance, by replacing 40 strands of 20 nucleotides by one strand of 800 nucleotides—increases the PR_25_+RNA critical temperature by zipping together large chunks of RNA that would otherwise be driven away by the dominant RNA– RNA electrostatic repulsion at physiological conditions. Thus, increasing the length of an RNA chain at constant nucleotide concentration, allows a higher density of PR_25_ bonds per RNA length, and an overall higher connected condensed liquid. A similar behaviour has been experimentally observed for the P-granule protein PGL-3, which has limited ability to undergo LLPS in absence of RNA. However, in presence of long (*>*600-mer) RNA strands, its phase separation is largely enhanced [122]. Also consistent with our observations, enrichment of long mRNA in stress granules [28, 123, 124] and NEAT1 RNA (∼23000-mer non-coding RNA transcripts) in paraspeckles [125, 126] promotes phase-separation of such membraneless organelles.

### C. RNA length has distinct effects on the stability of condensates driven by homotypic *versus* heterotypic interactions

To test the universality of these observations, we now employ our minimal protein model [67, 95–97], in which proteins are represented by patchy colloids [88] and RNA as a Lennard-Jones chain. This allows us to go beyond protein sequence, and assess the thermodynamic parameters that explain the general differences between the impact of RNA length on homotypic phase separation *versus* RNA–protein complex coacervation.

We start by computing the phase diagram of a minimal scaffold protein that, like FUS, is able to phase separate on its own via homotypic interactions. The scaffold protein is represented by a patchy particle decorated with 3-binding sites in a planar arrangement separated by 120 degrees angles (Figure 1 (a) Right). As shown in Ref. [67], below a reduced temperature of *T* ^*\**^ = 0.09 (see details on reduced units in the Supplementary Material), the scaffold proteins undergo phase separation (black curve of Figure 4 (a)). Crucially, when adding a self-avoiding flexible polymer that mimics RNA, we qualitatively recapitulate the impact on phase behaviour that we observed for FUS (Figure 2 (a)) with our residue-resolution coarse-grained simulations. That is, adding a moderate concentration of RNA, increases the critical temperature modestly (by about ∼35%), but changing the length of RNA (while keeping the nucleotide concentration constant) has a marginal effect (Figure 4 (a)).

**Figure 4:**
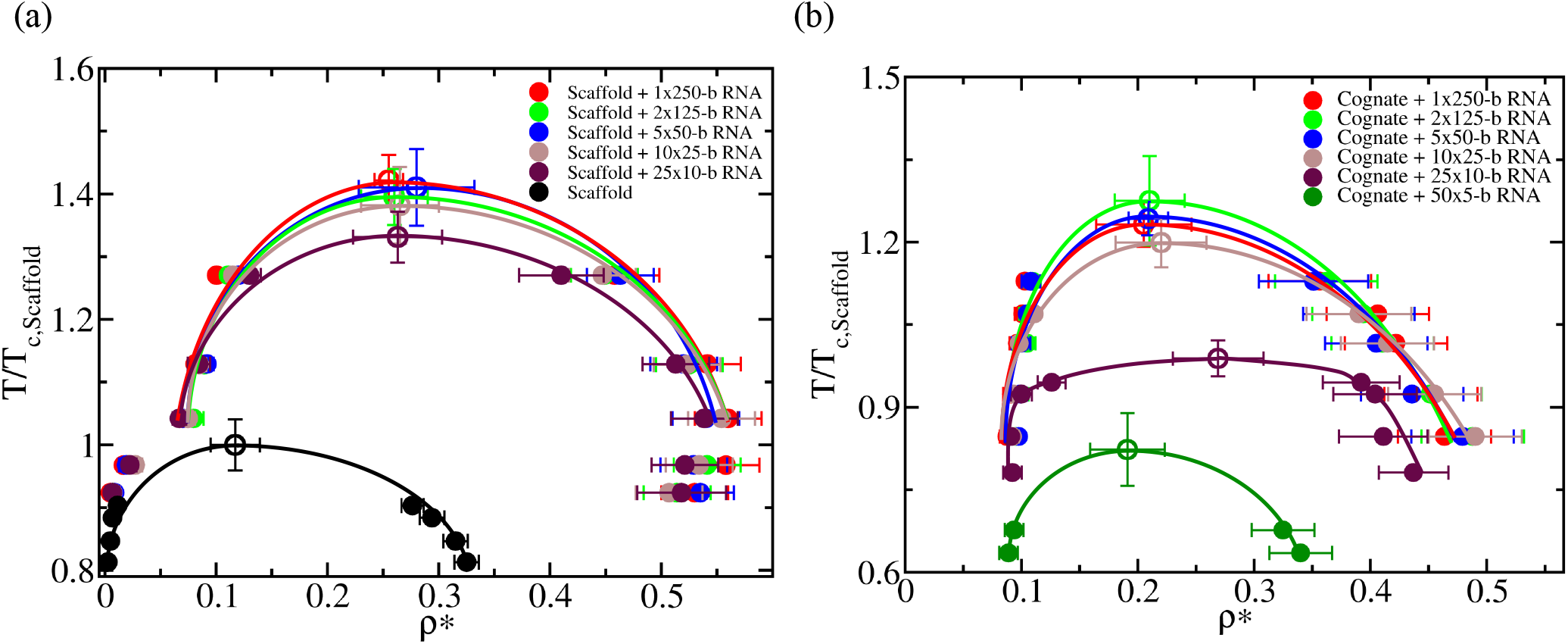
(a) Phase diagram in the temperature–density plane for a scaffold protein that, like FUS, can phase separate via homotypic protein interactions (black curve), and for mixtures with a fixed RNA/protein concentration using different RNA strand lengths as indicated in the legend. (b) Phase diagram in the temperature-density plane for a cognate protein that, like PR_25_, does not exhibit LLPS on its own, and that only undergoes LLPS upon addition of RNA strands. The RNA concentration in both panels was kept constant in all simulations at a 0.25 nucleotide/protein ratio. Filled circles represent the coexisting densities evaluated from DC simulations, while empty circles depict the critical temperatures estimated from the law of rectilinear diameters and critical exponents near the critical temperature [116]. Temperature in both panels has been normalized by the critical temperature of the pure scaffold system, 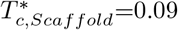 in reduced units (empty black circle in (a)).

Now we focus on the phase behaviour of a minimal cognate protein that, like PR_25_, cannot phase separate on its own (Figure 4 (b)). Our cognate proteins are represented by patchy particles with 2-binding sites in a polar arrangement, which has been shown to form linear chains [84, 88] instead of the well-connected percolated networks needed for sustaining condensates [97]. For the minimal cognate proteins, we obtain a phase behavior similar to that of PR_25_; when increasing the length of RNA (while keeping the nucleotide/protein ratio constant), the critical temperature of the mixture considerably increases (Figure 4 (b)). However, after reaching a certain RNA length that is much longer that the size of the proteins (i.e., ∼ 50 times longer, which in this minimal model can be tested) [48], the LLPS enhancement plateaus.

Next we analyze the density of protein–protein and protein–RNA contacts as a function of RNA length (Figure 5 (a) and (b)), to further elucidate the origins of the distinct behavior for scaffold and cognate proteins. We observe a similar trend in terms of the predicted liquid-network connectivity with our minimal model as that found using sequence-dependent coarse-grained simulations (Figure 3 (a) and (b)). When LLPS is mainly driven by homotypic scaffold–scaffold interactions, scaffold–scaffold and scaffold–RNA contacts remain roughly constant as the length of RNA increases. In contrast, when LLPS is significantly driven by RNA– protein (i.e., cognate protein) heterotypic interactions, the number of cognate–RNA contacts considerably augments with RNA length (until the RNA size is much larger than that of the proteins; Figure 5 (b)). While for the minimal scaffold proteins the increase in scaffold–scaffold and scaffold–RNA contacts with RNA length is smaller than a 5-10% (Figure 5 (a)), for cognate proteins such increase is higher than a factor of 3, which is a significant difference considering that in both cases RNA/protein ratios are kept constant. The variation in the critical temperature as a function of RNA length is also depicted in Figure 5 (c), where the consequences of the dissimilar liquid-network connectivity [97] that both type of proteins can establish to self-assemble – homotypic *vs*. heterotypic interactions – manifest.

**Figure 5:**
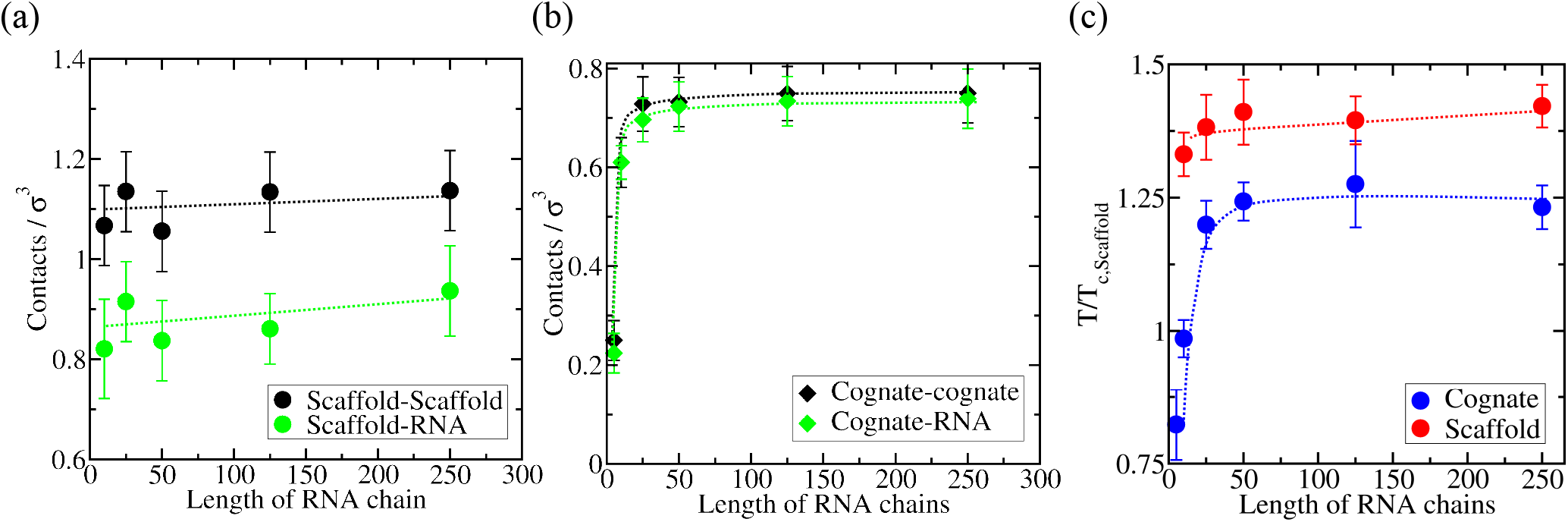
Density of LLPS-stabilizing contacts as a function of RNA length plotted separately for protein–protein contacts (black circles) and protein–RNA contacts (green circles) for a minimal RNA-binding scaffold protein model wherein scaffold proteins can phase separate via homotypic interactions (a) and an RNA-binding cognate protein model wherein cognate proteins can only phase separate via heterotypic RNA–protein interactions (b). Calculations are performed at 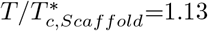 for the RNA/scaffold system and 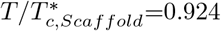 for the RNA/cognate protein system. The RNA/protein concentration was kept at a constant nucleotide/protein ratio of 0.25 in both cases. (c) Critical temperature *versus* RNA length plot for both mixtures, scaffold proteins + RNA (red) and cognate proteins + RNA (blue).

In agreement with the preceding results, Zacco *et al*. [25] found that longer RNA strands present weaker dissociation constants with N-RRM1-2 domains of TDP-43 (which, like PR_25_, cannot phase separate on their own at physiological conditions) than 3-fold shorter RNA strands. Moreover, another study by Maharana *et al*. [27] showed that smaller RNAs were more potent than larger ones in solubilizing protein condensates at high RNA concentration, which in turn, indirectly supports our observations that very short RNA strands can remotely promote LLPS for proteins that heavily rely on heterotypic interactions. Furthermore, besides controlling condensate stability, RNA has been suggested to play a critical role in regulating the dynamics of many membraneless organelles [21, 27, 30, 127, 128]. In that respect, Zhang *et al*. [129] showed that the RNA-binding protein Whi3 phase separates into liquid-like droplets wherein biophysical properties can be subtly tuned by changing the concentration and length of the mRNA binding partner, finding that larger RNA content increases Whi3 droplet viscosity. On the other hand, RNA has been shown to yield opposite effects in LAF-1 condensates when short strands (50 nt) were introduced [38]. Nonetheless, when long RNAs were used (up to 3,000 nt), LAF-1 condensates presented significantly higher viscosity [39]. Since the impact of RNA length and concentration in condensate density has been recently shown to be a good proxy of condensate dynamics (i.e., droplet viscosity and protein diffusion) [27, 39, 48], the reported variations in droplet density as a function of RNA length and temperature presented here in Figure 2 and Figure 4, can be also considered as good indicators of the impact that RNA length produces on RBP–RNA droplet transport properties. Therefore, RNA lengths that promote higher droplet density should also lead to important enhancements in droplet viscosity [48].

## III. CONCLUSIONS

Using a two-resolution simulation approach we demonstrate how variations in RNA length can yield non-trivial effects in the stability of RBP condensates. We find that in condensates sustained by homotypic protein–protein interactions, RNA behaves as a LLPS enhancer that subtly augments the stability of the condensates irrespective of its length, while, in condensates sustained by heterotypic protein–RNA interactions, RNA acts as a LLPS enabler that increases the stability of the condensates in a RNA-length-dependent manner.

Our findings for FUS and PR_25_ polyU systems using sequence-dependent coarse-grained simulations in parellel with our results for the miminal protein/RNA model suggest that when protein–protein LLPS-stabilising interactions are substantially higher than protein–RNA contacts, like in FUS or in our archetypal scaffold protein model, it is the RNA concentration rather than its chain length what critically modulates the condensate stability (at least for strands larger than 50–80 nucleotides or of comparable length to that of the proteins). Nevertheless, when protein–RNA intermolecular contacts contribute similarly or even higher than homotypic protein–protein interactions, like in PR_25_ peptides or in our minimal cognate–RNA model, not only the RNA concentration, but also the RNA chain length play a major role in controlling RBP condensate stability. Our study demonstrates how RNA participation in biological phase transitions is not uniform and should be carefully considered to the same extent as the wide amalgam of protein molecular interactions involved in the formation of biomolecular condensates.

## ACKNOWLEDGEMENTS

This project has received funding from the European Research Council (ERC) under the European Union Horizon 2020 research and innovation programme (grant agreement No 803326). J. R. E. acknowledges funding from the Oppenheimer Fellowship and from Emmanuel College Roger Ekins Research Fellowship. I. S. B. acknowledges funding from the Oppenheimer Fellowship, EPRSC Doctoral Programme Training number EP/T517847/1 and Derek Brewer Emmanuel College scholarship. J. A. J. is a Junior Research Fellow at Kings College. This work has been performed using resources provided by the Cambridge Tier-2 system operated by the University of Cambridge Research Computing Service (http://www.hpc.cam.ac.uk) funded by EPSRC Tier-2 capital grant EP/P020259/1.

