## Supplementary Material for "RNA length has a non-trivial effect in the stability of biomolecular condensates formed by RNA-binding proteins"

### I. MODELS AND SIMULATION DETAILS

#### A. Mpipi Model

For our simulations of FUS and $PR_{25}$ with poly-U, we employ the high-resolution sequence-dependent coarse-grained model Mpipi [1], which describes almost quantitatively the temperature-dependent LLPS phase behaviour of different protein condensates such as that of fused in sarcoma (FUS). In addition, this model correctly predicts the multiphase behaviour of the PolyR/PolyK/PolyU system, and recapitulates experimental LLPS trends for sequence mutations on FUS, DDX4 NTD and LAF-1 RRG domain variants [1]. Within this force field, electrostatic interactions are modelled with a Coulomb term with Debye–Huckel electrostatic screening [2], given by the sum over all particle-particle (i,j) interactions as:

$$E_{elec} = \sum_{i,j} \frac{q_i q_j}{4\pi\epsilon_r\epsilon_0 r_{ij}} exp(-\kappa r_{ij}) \tag{1}$$

where q is the charge, $\epsilon_r = 80$ is the relative dielectric constant of water, $\epsilon_0$ is the electric constant, $\kappa^{-1} = 795$ pm is the Debye screening length, and $r_{ij}$ is the distance separating particles $i$ and $j$. For these interactions, a Coulomb cut-off of 3.5 nm is employed. The non-bonded interactions between protein/RNA beads are modelled via the Wang–Frenkel potential [3]

$$E_{WF} = \sum_{i,j} \epsilon_{ij}\alpha_{ij} \left[\left(\frac{\sigma_{ij}}{r_{ij}}\right)^{2\mu_{ij}} - 1\right] \left[\left(\frac{\sigma_{ij}}{r_{ij}}\right)^{2\mu_{ij}} - 1\right]^{2\nu_{ij}} \tag{2}$$

where

$$\alpha_{ij} = 2\nu_{ij}\left(\frac{R_{ij}}{\sigma_{ij}}\right)^{2\mu_{ij}} \left[\frac{2\nu_{ij}+1}{2\nu_{ij}\left[\left(\frac{R_{ij}}{\sigma_{ij}}\right)^{2\mu_{ij}} - 1\right]}\right]^{2\mu_{ij}+1} \tag{3}$$

representing $\sigma$ the molecular diameter of each residue/nucleotide and $\epsilon$ the interaction strength between distinct amino acids and nucleotides ($i$ and $j$). $\mu_{ij}$ and $R_{ij}$ are constant model parameters set to $\mu_{ij} = 1$ and $R_{ij} = 3\sigma_{ij}$ for every interaction, while $\sigma_{ij}$ and $\epsilon_{ij}$ are specified for each pair of interaction in Ref. [1]. Finally, bond energy is computed with an harmonic bond potential of the following form:

$$E_{bond} = \sum_b \frac{1}{2}k(r_b - r_0) \tag{4}$$

where $b$ is the total number of bonds, $r_b$ is the bond distance, $k$=8.03 $\mathrm{Jmol^{-1}pm^{-2}}$ is the spring constant and $r_0$ is the bond reference position, set to 381 pm and 500 pm for protein and RNA bonds respectively. For further details on this force field and the full list of the model parameters please see Ref. [1].

#### B. Minimal protein/RNA model

For the minimal coarse-grained simulations, we employ a patchy particle model [4–7] in which proteins are described by a pseudo hard-sphere (PHS) potential [8] that accounts for their excluded volume:

$$E_{PHS} = \sum_{i<j} \begin{cases} \lambda_r (\frac{\lambda_r}{\lambda_a})^{\lambda_a} \varepsilon_R \left[ (\frac{\sigma}{r_{ij}})^{\lambda_r} - (\frac{\sigma}{r_{ij}})^{\lambda_a} \right] + \varepsilon_R; & \text{if } r < (\frac{\lambda_r}{\lambda_a})\sigma \\ 0; & \text{if } r \geq (\frac{\lambda_r}{\lambda_a})\sigma \end{cases}. \quad (5)$$

where $\lambda_a$ =49 and $\lambda_r$=50 are the exponents of the attractive and repulsive terms respectively, and $\varepsilon_R$ accounts for the energy shift of the pseudo hard-sphere. On top of this, we add a continuous square-well (CSW) potential for modeling the different protein binding sites, therefore mimicking protein multivalency:

$$E_{CSW} = \sum_{i<j} -\frac{1}{2}\epsilon_{CSW} \left[ 1 - tanh(\frac{r_{ij} - r_w}{\alpha}) \right] \quad (6)$$

where $\epsilon_{CSW}$ is the depth of the potential energy well, $r_w$ the radius of the attractive well, and $\alpha$ controls the steepness of the well. We choose $\alpha = 0.005\sigma$ and $r_w = 0.12\sigma$ so that each binding site can only interact with another single one. RNA-protein interactions are modeled with a standard Lennard-Jones (LJ) potential [9]:

$$E_{LJ} = \sum_{i<j} 4\epsilon_{LJ} \left[ \left(\frac{\sigma}{r_{ij}}\right)^{12} - \left(\frac{\sigma}{r_{ij}}\right)^{6} \right] \quad (7)$$

where $\epsilon_{LJ}$ measures the depth of potential and $\sigma$ the excluded volume between proteins and RNA. The Lennard-Jones potential is employed between proteins and RNA beads, while RNA-RNA interactions are modelled via the PHS potential [8]. In this way, we model RNA as a self-repulsive polymer of bonded hard spheres, via sites that are not the protein-protein binding sites, hence, if one protein is bonded to RNA, it can still bind to other proteins. The mass of each patch is a 5% of the central PHS particle mass, which is set to 3.32x$10^{-26}$kg, despite being this choice irrelevant for equilibrium simulations. This 5% ratio fixes the moment of inertia of the patchy particles (our minimal proteins). The molecular diameter of the proteins, both scaffold and cognate proteins, as well as the RNA beads is $\sigma$ = 0.3405 nm, and the value of $\varepsilon_R/k_B$ is 119.81K. With this model, we express magnitudes in reduced units: reduced temperature is defined as $T^* = k_BT/\epsilon_{CSW}$, reduced density as $\rho^* = (N/V)\sigma^3$, reduced pressure as $p^* = p\sigma^3/(k_BT)$, and reduced time as $\sqrt{\sigma^2 m/(k_BT)}$. In order to keep the PHS interaction as similar as possible to a pure HS interaction, we fix $k_BT/\varepsilon_R$ at a value of 1.5 as suggested in Ref. [8] (fixing $T = 179.71$K). We then control the effective strength of the binding protein attraction by varying $\epsilon_{CSW}$ such that the reduced temperature, $T^* = k_BT/\epsilon_{CSW}$, is of the order of $\mathcal{O}(0.1)$. The cutoff distance for the interactions in this model are $1.17\sigma$ for the PHS and CSW potentials and $5\sigma$ for the LJ interactions. The $\epsilon_{LJ}/k_B$ for LJ interactions is set to 152.5K.

This model has been proven to qualitatively reproduce the effect of protein valency in LLPS [5], the enhancement of RNA-mediated LLPS with RNA-binding proteins [10] or the size conservation of condensates through interfacial free energy reduction [7].

#### C. Simulation details

Our Direct Coexistence simulations are performed in the NVT ensemble (i.e. constant number of particles (N), volume (V) and temperature (T)), for which we use a Nosé–Hoover thermostat [11, 12] with a relaxation time of 5ps for the Mpipi model and 0.074 in reduced units for the patchy particle model. Since all our potentials are continuous and differentiable, we perform all our simulations using the LAMMPS Molecular Dynamics package[13]. Periodic boundary conditions are used in the three directions of space. The timestep chosen for the Verlet integration of the equations of motion is 10 fs for the Mpipi model and $3.7x10^4$ in reduced time units for the patchy particle model.

### II. COMPUTING PHASE DIAGRAMS VIA DIRECT COEXISTENCE

To calculate the coexisting densities of the phase diagrams, we employ the Direct Coexistence method [14–16]. Within this scheme, the two coexisting phases are simulated by preparing periodically extended slabs of the two phases, the condensed and the diluted one, in the same simulation box. We use an implicit solvent model; accordingly, the diluted phase (protein-poor liquid phase) and the condensed phase (protein-rich liquid phase) are effectively a vapour and a liquid phase, respectively. Once our DC simulations have reached equilibrium, we compute the density profile along the long axis of the box, and thus, we extract the density of the two coexisting phases (as shown in the Supporting Material of Ref. [10]). From the plateau of the condensed phase and the diluted one, we measure the density (avoiding the interfaces between both phases). To estimate the critical point of the phase diagrams, we use the universal scaling law of coexistence densities near a critical point [17], and the law of rectilinear diameters [18]:

$$\left(\rho_l^*(T^*) - \rho_v^*(T^*)\right)^{3.06} = d\left(1 - \frac{T*}{T_c^*}\right) \tag{8}$$

and

$$(\rho_l^* + \rho_v^*)/2 = \rho_c^* + s_2(T_c^* - T*) \tag{9}$$

where $\rho_l^*$, $\rho_v^*$ refer to the reduced coexisting densities of the condensed and diluted phases respectively, while $\rho_c^*$ is the critical reduced density, $T_c^*$ is the reduced critical temperature, and $d$ and $s_2$ are fitting parameters.

### III. RESULTS WITH THE HPS & KH MODELS FOR FUS AND PR$_{25}$

Additionally to the results presented in the main text of this article, we perform simulations of FUS and PR$_{25}$ in presence of RNA with different lengths at a constant RNA/protein concentration employing the hydrophobicity scale (HPS) model for the FUS-poly-U systems, and the Kim-Hummer (KH) model for those of PR$_{25}$ with poly-U. Both models are detailed in Ref. [19]. In these models, the solvent is also implicit as in the Mpipi [1]. The simulation details are the same provided for the Mpipi simulations in Section 1C. Here, poly-U RNA strands are mimicked as chains of glutamic acid as shown in Ref. [20]. In Figure 1 (a) we show how the phase behaviour of FUS and poly-U RNA deeply resembles the one observed in the main document using the Mpipi model (Figure 2 (a) of the main text): the critical temperature of FUS-RNA mixtures marginally depends on the length of the added poly-U strands. However, for PR$_{25}$-poly-U systems (Figure 1 (b)), there is a remarkable difference of a factor of 2 (almost 200K) between the shortest (20-nucleotide long RNA chains) and the longest ones (800-nucleotide long RNA chains) as observed in the main text for the Mpipi model (Figure 2 (b) of the main text).

Moreover, the same intermolecular contact analysis of Figure 3 of the main text was also performed for these simulations, providing similar results to those computed from the Mpipi model calculations. In Figure 2(a), it can be seen how FUS-FUS contacts are mostly responsible of holding the protein condensates (displaying about 4 times more contacts per nm$^3$ than the RNA-FUS contacts), while in the PR$_{25}$-RNA condensates, heterotypic PR$_{25}$-RNA interactions are predominant respect to those of PR$_{25}$-PR$_{25}$ (Figure 2 (b)). Lastly, the length dependent behaviour of the critical temperature for PR$_{25}$ condensates is held as in the case of the Mpipi model (main document, Figure 3(c)), while for FUS-poly-U systems the critical temperature remains constant almost independently of the RNA chain length.

### IV. COMPUTING THE NUMBER OF INTERMOLECULAR CONTACTS

In Figures 3 and 5 of the main document, we show the number of intermolecular contacts per unit of volume. To compute such magnitude, we consider two amino acids to be in contact under a distance of 7.7$\AA$, which is the average of all the possible $\sigma_{ij}$ (eq. 2) multiplied by a factor 1.2 (typical value in $\sigma$ at which the attractive LJ/Wang-Frenkel interactions become very mild). The RNA-protein interactions are considered as contacts when the distance between a nucleotide and an amino acid is lower than 8.95$\AA$, which is the average $\sigma_{ij}$ between uridine and any other aminoacid, multiplied also by a factor of 1.2. The number of total contacts is averaged throughout entire converged simulations of about 2000 ns. Then, the contacts density is obtained by dividing the total number of contacts by the condensate volume.

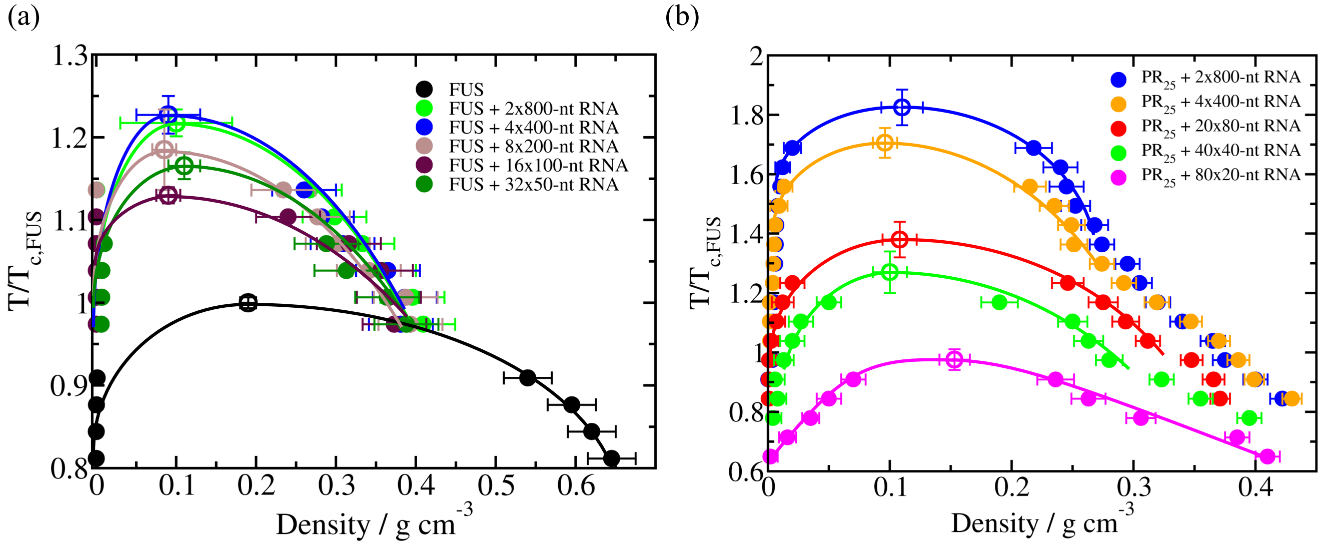


Figure 1: Temperature–density phase diagrams of FUS with Poly-U of different lengths at a constant PolyU/FUS mass ratio of 0.16, and for a pure system of FUS (black curve). (b) Temperature–density phase diagrams of $PR_{25}$ with RNA at different lengths at a constant RNA/$PR_{25}$ mass ratio of 0.57. In both (a) and (b) panels, filled circles represent the coexisting densities evaluated from DC simulations while empty circles depict the critical temperatures estimated from the law of rectilinear diameters and critical exponents [17] near the critical temperature. Temperature in both panels has been normalized by the critical temperature of pure FUS, $T_{c,FUS}$=309K (black empty circle in (a)).

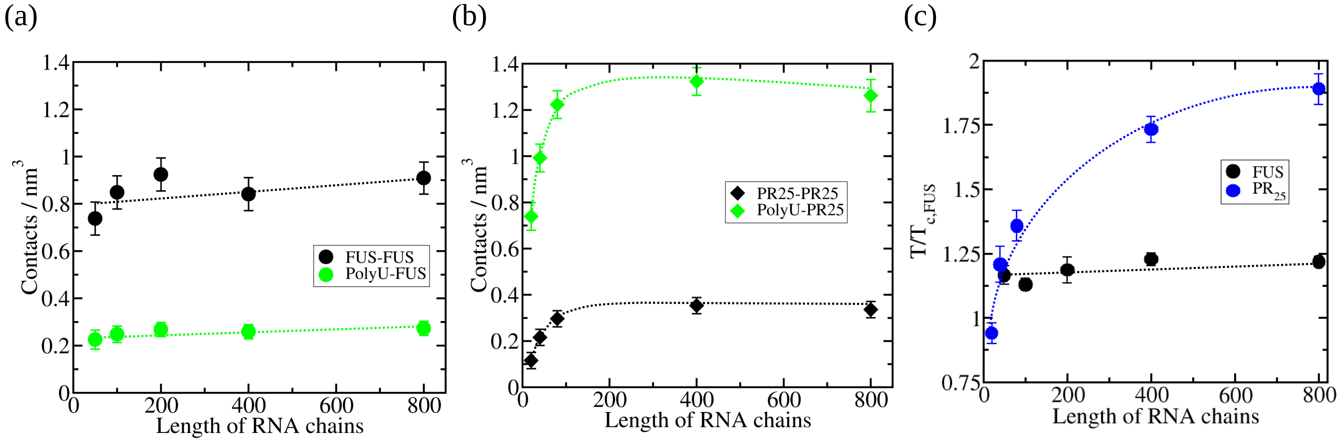


Figure 2: Density of LLPS-stabilizing intermolecular contacts within condensates as a function of RNA length plotted separately for protein–protein interactions (black circles) and protein–RNA interactions (green circles) for FUS–PolyU (a) and $PR_{25}$-PolyU mixtures (b). The temperature at which the intermolecular contacts were computed was $T/T_{\mathrm{c,FUS}}$=1.13 for FUS–RNA systems and $T/T_{\mathrm{c,FUS}}$=0.924 for $PR_{25}$–RNA mixtures (the highest temperature at which all systems with distinct RNA lengths can phase separate). (c) Critical temperature *versus* RNA length for FUS–RNA (red) and $PR_{25}$–RNA (blue) systems.

[1] A. J. Joseph, A. Reinhardt, A. Aguirre, P. Y. Chew, K. O. Russell, J. R. Espinosa, A. Garaizar, and R. Collepardo-Guevara, Nat. Comput. Sci. , in press (2021).

[2] P. Debye and E. Hückel, Physikalische Zeitschrift **24**, 185 (1923).
[3] X. Wang, S. Ramírez-Hinestrosa, J. Dobnikar, and D. Frenkel, Physical Chemistry Chemical Physics **22**, 10624 (2020).
[4] J. R. Espinosa, A. Garaizar, C. Vega, D. Frenkel, and R. Collepardo-Guevara, J. Chem. Phys **150**, 224510 (2019).
[5] J. R. Espinosa, J. A. Joseph, I. Sanchez-Burgos, A. Garaizar, D. Frenkel, and R. Collepardo-Guevara, Proceedings of the National Academy of Sciences (2020).
[6] I. Sanchez-Burgos, J. R. Espinosa, J. A. Joseph, and R. Collepardo-Guevara, Biomolecules **11**, 278 (2021).
[7] I. Sanchez-Burgos, J. A. Joseph, R. Collepardo-Guevara, and J. R. Espinosa, bioRxiv (2021).
[8] J. Jover, A. J. Haslam, A. Galindo, G. Jackson, and E. A. Müller, Journal of Chemical Physics **137** (2012), 10.1063/1.4754275.
[9] J. E. Jones, Proceedings of the Royal Society of London. Series A, Containing Papers of a Mathematical and Physical Character **106**, 441 (1924).
[10] J. A. Joseph, J. R. Espinosa, I. Sanchez-Burgos, A. Garaizar, D. Frenkel, and R. Collepardo-Guevara, Biophysical Journal **120**, 1219 (2021).
[11] S. Nosé, The Journal of Chemical Physics **81**, 511 (1984).
[12] W. G. Hoover, Phys. Rev. A **31**, 1695 (1985).
[13] S. Plimpton, Journal of Computational Physics **117**, 1 (1995).
[14] A. J. Ladd and L. V. Woodcock, Chemical Physics Letters **51**, 155 (1977).
[15] R. García Fernández, J. L. F. Abascal, and C. Vega, The Journal of Chemical Physics **124**, 144506 (2006).
[16] J. R. Espinosa, E. Sanz, C. Valeriani, and C. Vega, Journal of Chemical Physics **139** (2013), 10.1063/1.4823499.
[17] J. S. Rowlinson and B. Widom, *Molecular theory of capillarity* (Courier Corporation, 2013).
[18] J. A. Zollweg and G. W. Mulholland, The Journal of Chemical Physics **57**, 1021 (1972).
[19] G. L. Dignon, W. Zheng, Y. C. Kim, R. B. Best, and J. Mittal, PLoS Computational Biology **14** (2018), 10.1371/journal.pcbi.1005941.
[20] G. Krainer, T. J. Welsh, J. A. Joseph, J. R. Espinosa, S. Wittmann, E. de Csilléry, A. Sridhar, Z. Toprakcioglu, G. Gudiškytė, M. A. Czekalska, *et al.*, Nature Communications **12**, 1 (2021).
